# Amphetamine disrupts haemodynamic correlates of prediction errors in nucleus accumbens and orbitofrontal cortex

**DOI:** 10.1101/802488

**Authors:** Emilie Werlen, Soon-Lim Shin, Francois Gastambide, Jennifer Francois, Mark D Tricklebank, Hugh M Marston, John R Huxter, Gary Gilmour, Mark E Walton

**Author notes:** deceased (Soon-Lim Shin).

## Abstract

In an uncertain world, the ability to predict and update the relationships between environmental cues and outcomes is a fundamental element of adaptive behaviour. This type of learning is typically thought to depend on prediction error, the difference between expected and experienced events, and in the reward domain this has been closely linked to mesolimbic dopamine. There is also increasing behavioural and neuroimaging evidence that disruption to this process may be a cross-diagnostic feature of several neuropsychiatric and neurological disorders in which dopamine is dysregulated. However, the precise relationship between haemodynamic measures, dopamine and reward-guided learning remains unclear. To help address this issue, we used a translational technique, oxygen amperometry, to record haemodynamic signals in the nucleus accumbens (NAc) and orbitofrontal cortex (OFC) while freely-moving rats performed a probabilistic Pavlovian learning task. Using a model-based analysis approach to account for individual variations in learning, we found that the oxygen signal in the NAc correlated with a reward prediction error, whereas in the OFC it correlated with an unsigned prediction error or salience signal. Furthermore, an acute dose of amphetamine, creating a hyperdopaminergic state, disrupted rats’ ability to discriminate between cues associated with either a high or a low probability of reward and concomitantly corrupted prediction error signalling. These results demonstrate parallel but distinct prediction error signals in NAc and OFC during learning, both of which are affected by psychostimulant administration. Furthermore, they establish the viability of tracking and manipulating haemodynamic signatures of reward-guided learning observed in human fMRI studies using a proxy signal for BOLD in a freely behaving rodent.

## Introduction

The world is an uncertain place, where behaviour of animals must continuously change to promote optimal survival. Learning to predict the relationship between environmental cues and significant events is a critical element of adaptive behaviour. It is hypothesised that adaptive behaviour depends upon comparisons of neural representations of cue-evoked expectations of events with events that actually occurred. Mismatch between these two representations is defined as a prediction error, and is likely a vital substrate by which accuracy of ensuing predictions about cue-event relationships can be improved. Prediction errors related to receipt of reward have been strongly associated with dopaminergic neurons and their projections to frontostriatal circuits [1–4]. In rodents and humans, presentation of reward-predicting cues cause an increase in dopaminergic neuron activity and dopamine release in terminal regions, not only in proportion to the expected value of the upcoming reward but also to the deviation from that expectation when the reward is actually delivered [5–12]. Furthermore, experimental disruption of dopaminergic transmission can impair formation of appropriate cue-reward associations [13–15].

From a human perspective, elements of reward learning can be disrupted in a variety of neuropsychiatric conditions where dopaminergic dysfunction may play a central role [16–19]. For example, patients with major depressive disorder or schizophrenia can be insensitive to reward and display impairments in reward learning behaviours [20–24]. Neuroimaging studies suggest that activation of parts of ventral striatum and frontal cortex, or changes in functional connectivity with these regions, may be an important neurophysiological correlate of reward learning impairments [24–29]. However, not all studies show the same patterns of changes [e.g., 30] and there is still uncertainty over whether the blunting of neural responses reflects a primary aetiology in these disorders. While the potential links between dopaminergic dysregulation, disrupted neural signatures of reward-guided learning, and neuropsychiatric symptoms are manifest, strong direct evidence is currently lacking.

To help bridge this gap, we used constant-potential amperometry to monitor haemodynamic responses simultaneously in the nucleus accumbens (NAc) and orbitofrontal cortex (OFC) in rats performing a reward-driven, probabilistic Pavlovian learning task. NAc and OFC regions both receive dopaminergic input, and have been previously implicated in representing the expected value of a cue to guide reward learning behaviour [31,32]. Amperometric tissue oxygen [T_O2_] signals likely originate from equivalent physiological mechanisms as fMRI BOLD signals [33–35], allowing cross-species comparisons of behaviourally driven haemodynamic signals in awake animals. The aim of this study was to define amperometric signatures of cue-evoked expectation of reward and prediction error in these regions and then investigate how they are modulated by administration of amphetamine.

Amphetamine is known to modify physiological dopamine signalling [36] and, in humans, even a single dose of a stimulant like methamphetamine can cause an increase in mild psychotic symptoms [37,38]. While amphetamine can promote behavioural approach to rewarded cues [39], it can also impair conditional discrimination performance [40] and impair the influence of probabilistic cue-reward associations on subsequent decision making [41]. We hypothesised that amphetamine would disrupt discriminative responses to cues during performance of a probabilistic Pavlovian task, with concomitant changes to the haemodynamic correlates of reward expectation and prediction error in the NAc and OFC.

## Methods

See SI for detailed methods.

### Animals

All experiments were conducted in accordance with the United Kingdom Animals (Scientific Procedures) Act 1986. Adult male Sprague Dawley rats (Charles River, UK) were used in the present studies (*n*=36). 4 animals did not contribute to the behavioural dataset owing to poor O_2_ calibration responses and the data from an additional 2 animals could not be included owing to a computer error. During testing, they were maintained at no less than 85% of their free-feeding weight relative to their normal growth curve. Prior to the start of any training or testing, all animals underwent surgical procedures under general anaesthesia to implant carbon paste electrodes targeted bilaterally at the NAc and OFC.

### O2 amperometry data recording

O2 signals were recorded from the NAc and OFC using constant potential amperometry (−650 mV applied for the duration of the session) as described previously [35,42].

### Probabilistic Pavlovian conditioning task

The task was a probabilistic Pavlovian learning task performed in standard operant chambers. Each trial consisted of a 10s presentation of one of two auditory cues (3 kHz pure tone at 77dB or 100 Hz clicker at 76 dB) followed immediately by either delivery or omission of reward (4 × 45 mg sucrose food pellets). One of the auditory cues (CS_High_) was followed by reward delivery on 75% of trials, the other (CS_Low_) was rewarded on 25% of trials. Each session consisted of a total of 56 cue presentations, with an average inter-trial interval of 45s (range 30-60s). Standard training took place over 9 sessions and session 10 consisted of the drug challenge (see Figure S1).

### Pharmacological manipulations

D-amphetamine sulphate (Sigma,UK) was dissolved in 5% (w/v) glucose solution, and pH adjusted towards neutral with the dropwise addition of 1M NaOH as necessary. Amphetamine was dosed at 1mg/kg (free weight) via the intraperitoneal route.

### Behavioural modelling

Head entries during the 10 seconds cue presentation were modelled using variations of a Rescorla-Wagner model (Rescorla & Wagner 1972). We started with a model with a single free parameter, the learning rate α, and compared this against other models that also included free parameters specifying: (a) cue-specific learning rates (i.e., a cue salience term, β); (b) separate learning rates for rewarded α_pos_ and non-rewarded trials α_neg_; and either (c) cue-independent *k* or (d) cue-specific unconditioned magazine responding, *k*_*Clicker*_ and *k*_*Tone*_. To capture additional trial-by-trial variance, we also included either trial-specific or recency-weighted pre-cue response rates. To compare the models, we used the Bayesian information criterion (BIC), which penalises the likelihood of a model by the number of parameters and the natural logarithm of the number of data points.

### Data analysis

#### Behaviour

We analysed the average number of head entries into the food magazine during presentation during either the CS_High_ or CS_Low_ cues during the 9 days of training and then during the pre-drug day (day 9 of training) with the drug challenge day.

#### Amperometry

We performed two sets of complementary analyses: (i) model free analyses, where we investigated the average signals in NAc and OFC during cue presentation or in the 30s after outcome delivery over the course of learning and after amphetamine administration, and (ii) model based analyses where we regressed the same signals against estimates from our computational model.

## Results

### Behavioural performance during probabilistic learning

We trained rats on a two-cue probabilistic Pavlovian learning paradigm. One cue – CS_High_ – was associated with reward delivery on 75% of trials and the other – CS_Low_ – on 25% of trials (Figure 1A). As can be observed, animals learned to discriminate between the cues, increasingly making magazine responses during presentation of the CS_High_ but showing little change in behaviour upon presentation of CS_Low_ as training progressed (main effect of CS: F_1,27_=39.92, p<0.001; CS × day interaction: F_2.63,70.92_=6.12, p=0.001) (Figure 1B, Figure S2). Unexpectedly, however, there was also a substantial and consistent influence of the counterbalancing assignment on responding (CS × cue identity interaction: F_1,27_=48.17, p<0.001). Specifically, follow up pairwise comparisons showed that the animals in Group 1, where CS_High_ was assigned to be the clicker cue and CS_Low_ the pure tone (“CL1-T2”) exhibited strong discrimination between the cues throughout training (p<0.001; Figure 1C). By contrast, rats in Group 2 with the opposite CS – auditory cue assignment (“T1-CL2”), did not show differential responding to the cues in spite of the different reward associations (p=0.66).

**Figure 1.**
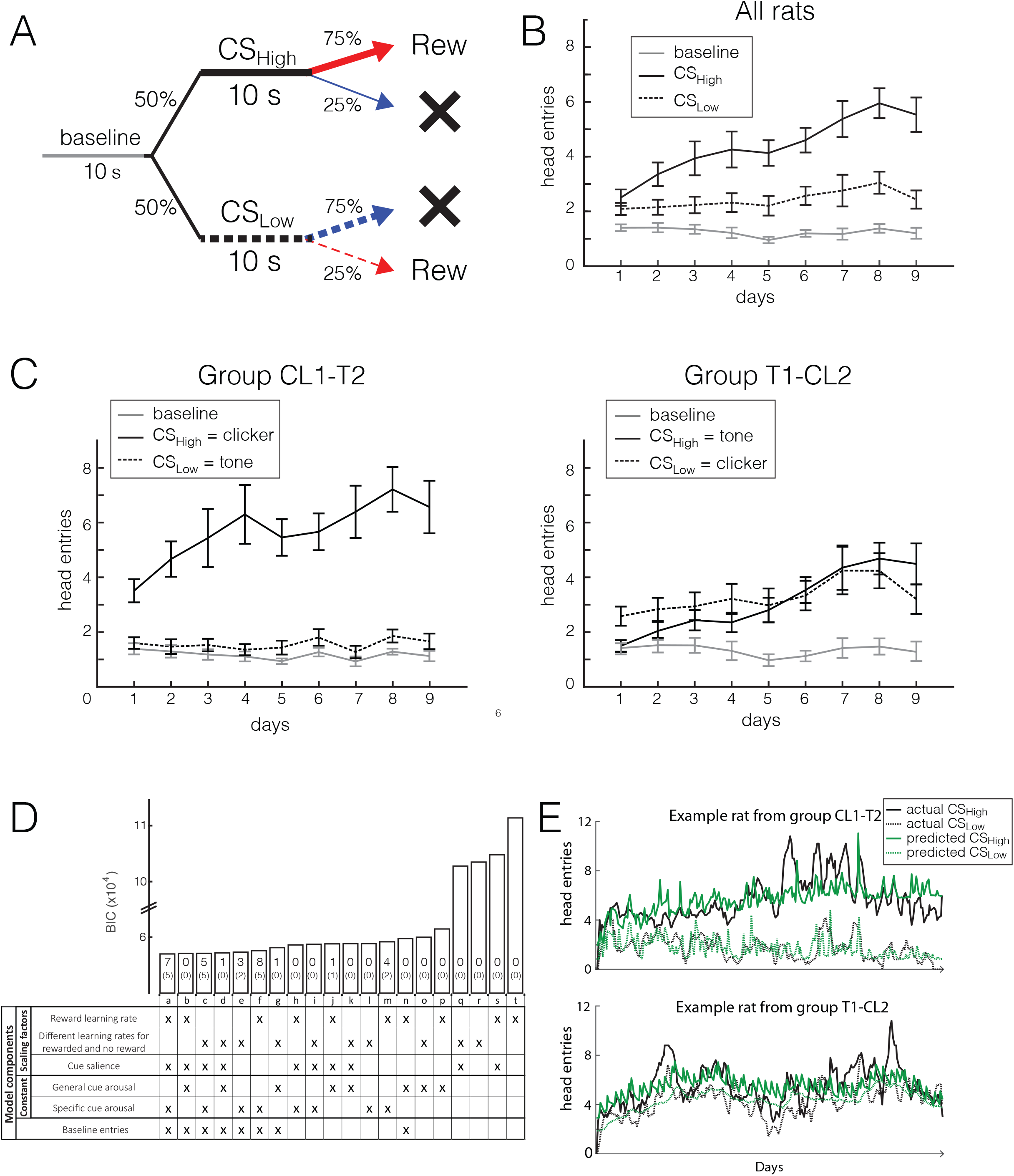
Task, behavioural performance and modelling. **A**: Schematic of the Pavlovian task. **B, C**: Average head entries (mean ± SEM) to the magazine during presentation of each cue or during the pre-Cue baseline period across the 9 sessions (Panel B, all animals; Panel C, Group C1-T2 only, where the CS_High_ was the clicker and CS_Low_ was the tone; Panel D, Group T1-C2 only, where the CS_High_ was the tone and CS_Low_ the clicker). **D**: Bayesian information criterion (BIC, a measure of the goodness of fit of the model) estimates for different learning models. The BIC penalises the likelihood of a model by the number of parameters and the natural logarithm of the number of data points. The model with the lowest BIC score was deemed to give a better fit of the data. Note, however, the patterns of results in the model-based analyses of amperometric signals remained unchanged if we used any of the 3 models that fitted best for a number of individual rats (models a, c, f). An ‘x’ in the table indicates the presence of the given component in the model. Numbers within each bar indicate the number of animals for which the given model had lowest BIC. **E**: Example of the model fits for two animals from the two counterbalancing groups.

To better understand how the cue identity was influencing this pattern of responding, we formally analysed how well different simple reinforcement learning models could describe individual rats’ Pavlovian behaviour. The preferred model included a cue salience parameter and a cue-specific unconditioned magazine responding term, as well as recency-weighted pre-cue responding parameter (Figure 1D). In particular, the constant term attributable to unconditioned cue-elicited magazine responding was higher on clicker than tone trials (Z=3.98, p<0.001, Wilcoxon signed ranks test; p<0.015 for Group 1 or 2 analysed separately). Therefore, once the difference in cue attributes was accounted for, rats’ behaviour could be well explained using this modified simple reinforcement learning model (Figure 1E).

### Both NAc and OFC haemodynamic signals track Pavlovian responding

We examined how T_O2_ responses in NAc and OFC (Figure 2) tracked animals’ learning of the appetitive associations and violations of their expectations. After exclusions for misplaced electrodes and poor quality of signals (see Supplementary Methods, Figure S3), 40 electrodes in 20 rats were included for analysis (NAc=25 electrodes from 15 rats, OFC=15 electrodes from 11 rats).

**Figure 2.**
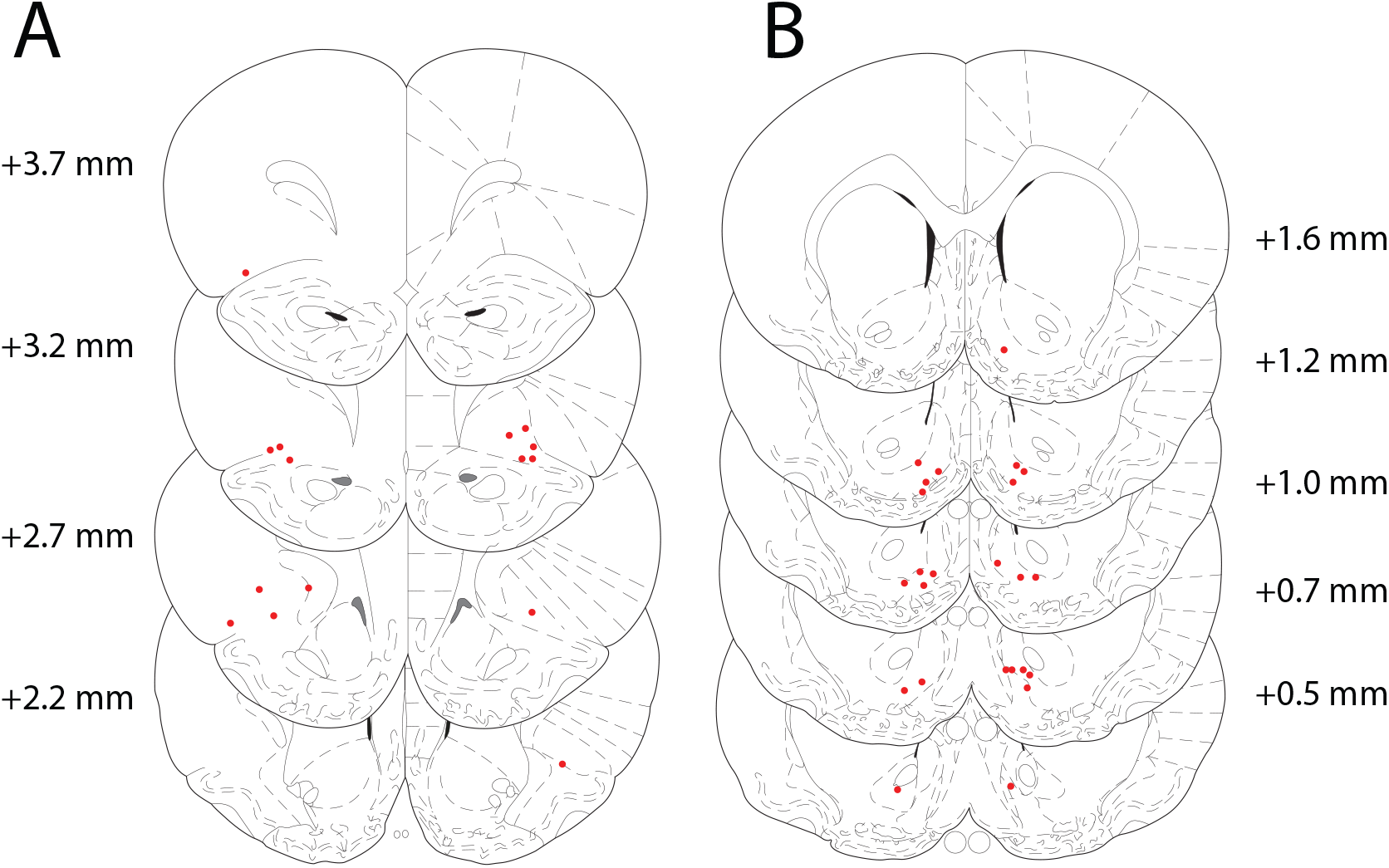
Electrode location in OFC (A) and NAc (B). The NAc electrodes were clustered mainly in the ventromedial NAc, including ventromedial core and shell, while the OFC electrodes were in the ventral orbital sector

We initially performed model free analyses to investigate the T_O2_ signals in response to presentation of the CS_High_ and CS_Low_ cues as the rats learned the reward associations.

T_O2_ responses during presentation of the cues changed markedly over training in a similar manner in both brain regions (main effects of cue and session: both F>9.49, p<0.001) (Figure 3A,B). In fact, analysis of the subset of animals with functional electrodes recorded simultaneously in NAc and OFC (n = 6 rats) showed a significant positive correlation between the signals recorded in each area (r^2^=0.41, p<0.01). Moreover, mirroring the behavioural data, the patterns of responses differed substantially according to the cue identity (CL1-T2 or T1-CL2). While the T_O2_ response following the CS_High_ developed similarly in both groups, there was a substantial difference in the CS_Low_ response, with average signals when the clicker was the CS_Low_ being significantly higher than when the tone was the CS_Low_ (Cue × cue identity interaction: F_1,36_=18.67, p<0.001; CL1-T2 v T1-CL2, CS_High_: p=0.32, CS_Low_: p<0.001).

**Figure 3.**
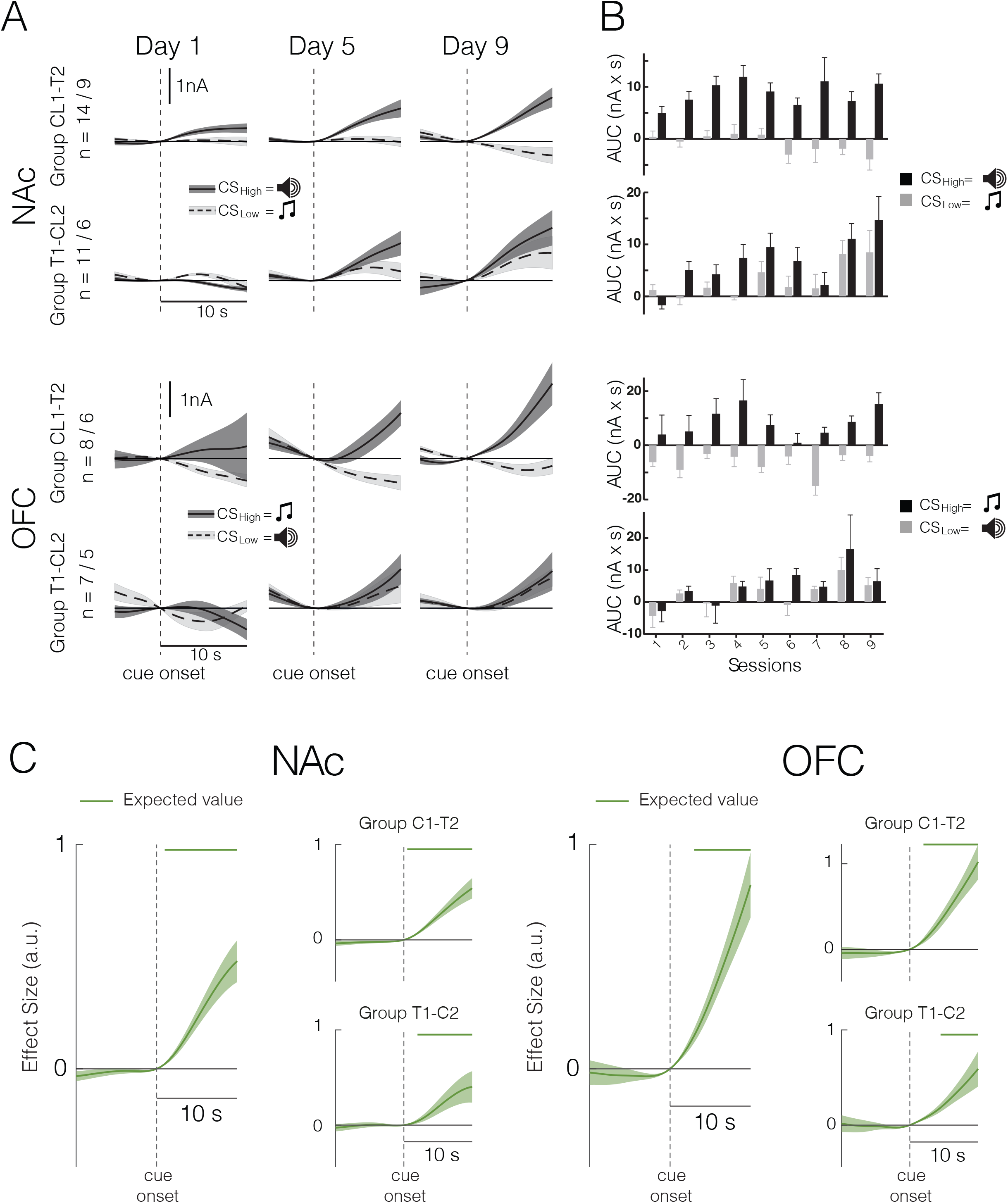
Haemodynamic correlates during cue presentation. **A**. T_O2_ responses on 3 sample days time-locked to cue presentation in the two counterbalance groups recorded from either NAc (upper panels) or OFC (lower panels). **B**. Average area-under-the-curve responses (mean ± SEM) extracted from 5-10 sec after cue onset for each cue across the 9 sessions. **C**. Average effect sizes in NAc (left panel) and OFC (right panel) from a general linear model relating T_O2_ responses to trial-by-trial estimates of the expected value associated with each cue. Main plots include all animals, insets show the analyses divided up into the two cue identity groups.

To establish the relationship between development of magazine responding and the T_O2_ signals, we regressed the model-derived estimates of cue value, *V*(*t*), against the trial-by-trial T_O2_ responses and found a significant positive relationship in both regions in both groups (Figure 3C). This was not simply a correlate of invigorated responding as cue value was a significantly better predictor than trial-by-trial magazine head entries (Figure S4). Therefore, once differences in cue identity are accounted for, it is possible to demonstrate that T_O2_ signals in both NAc and OFC track the expected value associated with each cue.

### Separate haemodynamic signatures of signed and unsigned prediction errors in NAc and OFC

We next investigated how the probabilistic delivery or omission of reward shaped T_O2_ responses in NAc and OFC, and how these signals were shaped by cue-elicited reward expectations as learning progressed. As there were significant interactions between brain region with cue and training stage and their combination (all F>3.76, p<0.028), we here analysed responses in the two regions separately.

We again first performed model free analyses, focusing on how the average outcome-evoked changes in T_O2_ responses were influenced by the preceding cue and how these adapted over training. As can be seen in Figure 4, the primary determinant of the signal change in the NAc was whether a reward was received (main effect of reward: F_1,36_=47.72, p<0.001). However, the size of reward and no reward signals, normalised to the time of outcome, depended on which cue had preceded the outcome and these patterns altered as learning progressed (significant cue × outcome and cue × outcome × training stage interactions, both F>18.669, p<0.001), suggesting a strong influence of expectation on T_O2_ responses. Follow up comparisons showed that there was a reduction in reward-elicited T_O2_ responses on CS_High_ trials as training progressed (CS_High_ rew, stage 1 v stage 3: p=0.016; stage 2 v stage 3: p=0.065). Unexpectedly, there was also a diminution of omission-elicited reductions in T_O2_ responses on these trials (CS_High_ no reward, stage 1 v stage 3: p=0.014), which, from Figure 4B, can be seen to be particularly prominent in the CL1-T2 group. By contrast, on CS_Low_ trials there was no meaningful change in T_O2_ responses to delivery or omission of reward throughout training (all p>0.37). As might be expected given its effect on behaviour and cue-elicited neural signals, cue identity again influenced outcome signals, resulting in a 4-way interaction of all the factors, as well as 2-way interactions between cue × identity and training stage × identity (all F>4.33, p<0.019). Importantly, however, when analysed separately, both counterbalance groups showed the key cue × outcome × training stage interaction (F>4.84, p<0.019).

**Figure 4.**
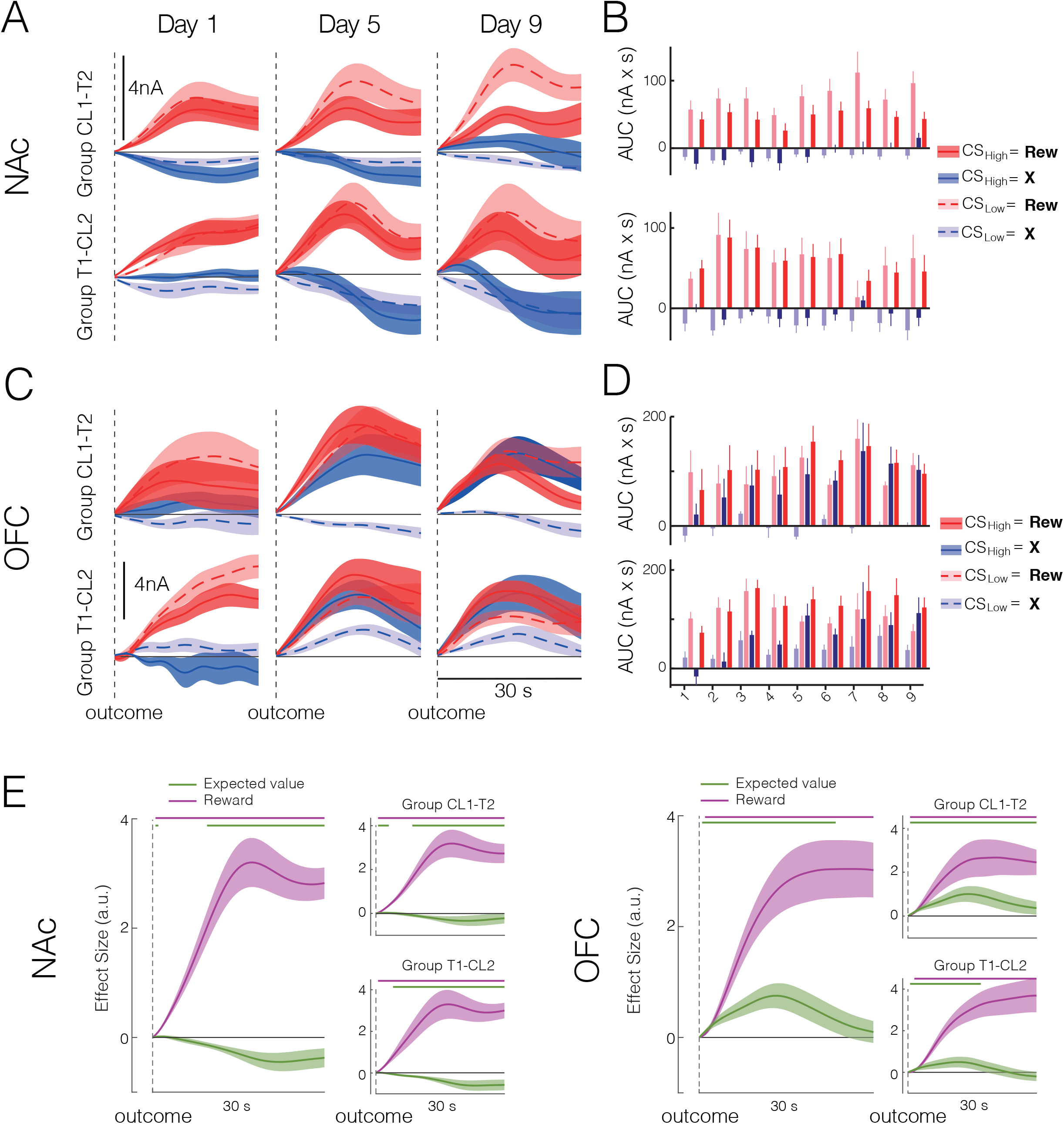
Haemodynamic correlates following outcome presentation. **A, C**. T_O2_ responses on 3 sample days, time-locked to outcome presentation (reward or no reward) after each cue in the two cue identity groups recorded from either NAc (panel A) or OFC (panel C). **B, D**. Average area-under-the-curve responses (mean ± SEM) extracted from 30 sec following the outcome after each cue across the 9 sessions (NAc panel A, OFC panel C). **E.** Average effect sizes in NAc (left panel) or OFC (right panel) from a general linear model relating T_O2_ responses to trial-by-trial estimates of the expected value associated with each cue and trial outcome (reward or no reward). Main plots include all animals, insets show the analyses divided into the two cue identity groups.

In the OFC, outcome was also a strong influence on T_O2_ responses (F_1,13_=51.18, p<0.001) and this was again shaped by the preceding cue (cue × outcome interaction: F_1,13_=5.37, p=0.037). However, unlike in NAc, there came to be an increasingly strong T_O2_ response when reward was *omitted*, particularly after CS_High_ (Figure 4C,D). This resulted in a cue × training stage interaction (F_1,13_=5.37, p=0.037; CS_High_ v CS_Low_, p=0.51 for the early training stage, p<0.001 for mid and late stages). While there were qualitative differences in responses in the two counterbalance groups, none of the interactions with this factor or the main effect reached significance (all p>0.069).

While these model-free analyses illustrate that the T_O2_ responses changed dynamically and differently over the course of training in the two brain regions, they do not clearly show whether either response might encode a teaching signal useful for learning, such as a reward PE: δ(*t*) = *V*(*t*) + *r*(*t*) − *V*(*t* − 1). Therefore, we next used model-based analyses to examine whether there was a relationship between T_O2_ responses across all sessions and the fundamental components of a reward PE: (i) a *positive* influence of outcome, *r*(*t*), and (ii) a *negative* influence of model-derived cue value, −*V*(*t* − 1). While both NAc and OFC T_O2_ responses showed a strong positive influence of outcome, only the NAc signals fulfilled both criteria of a reward PE by also exhibiting a significant negative influence of cue value; in OFC, by contrast, the cue value effect was positive (Figure 4E). We also examined whether reward PE-like T_O2_ responses in NAc were present throughout training. This showed that, while correlates of both NAc positive and negative reward PEs can be observed in rats that are still learning the cue-reward associations, once appropriately established only positive reward PEs remain evident (Figure S5).

Although the OFC T_O2_ responses do not correspond to a reward PE, the patterns of signals nonetheless still dynamically change over learning. As previous work has suggested that OFC neurons may signal the salience of the outcome for learning, we examined whether T_O2_ responses instead correlated with how unexpected each outcome was, corresponding to an *unsigned* PE. This analysis showed that each animal’s trial-by-trial unsigned PE had a strong positive influence on OFC signals (Figure S6). Again, this was present in both counterbalance groups.

Therefore, while the changes in NAc T_O2_ responses reflect how much better or worse an outcome was than expected, OFC T_O2_ responses indicate how surprising either was.

### Amphetamine disrupts cue-specific value encoding and prediction errors

Having established haemodynamic PE correlates in NAc and OFC, we next wanted to investigate how an acute dose of amphetamine (1 mg/kg), an indirect sympathomimetic known to potentiate dopamine release, influenced cue value and prediction error T_O2_ responses.

We first analysed baseline magazine responding in the pre-drug and the drug administration sessions. Although amphetamine caused a numeric increase in baseline responding, this was variable between animals − 7/15 rats given amphetamine showing a substantial increase in baseline magazine response rates, whereas the other 8/15 animals showed a decrease in response rates – and the drug × session interaction did not reach significance (F_1,26_=3.71, p=0.065). By contrast, there was a substantial and consistent change in cue-elicited responses (cue × session × drug interaction: F_1,26_=16.31, p<0.001). This was not caused by differences between the drug groups on the pre-drug session (no main effect or interaction with drug group: all F < 1.21, p > 0.28). Instead, as can be observed in Figure 5A, while both the vehicle and amphetamine groups responded more on average to the CS_High_ than the CS_Low_ on the pre-drug day (p<0.003), this discrimination was abolished after administration of the drug (CS_High_ v CS_Low_: p=0.35), but not the vehicle (p<0.001). Note that while there were again some differences between the counterbalance groups (cue × session × drug × identity interaction: F_1,26_=6.30, p=0.019), the effects of amphetamine administration were comparable in both groups (Amphetamine group: significant cue × session interaction, F_1,13_=9.047, p=0.01; no significant cue × session × identity interaction, F_1,13_=2.71, p=0.12).

**Figure 5.**
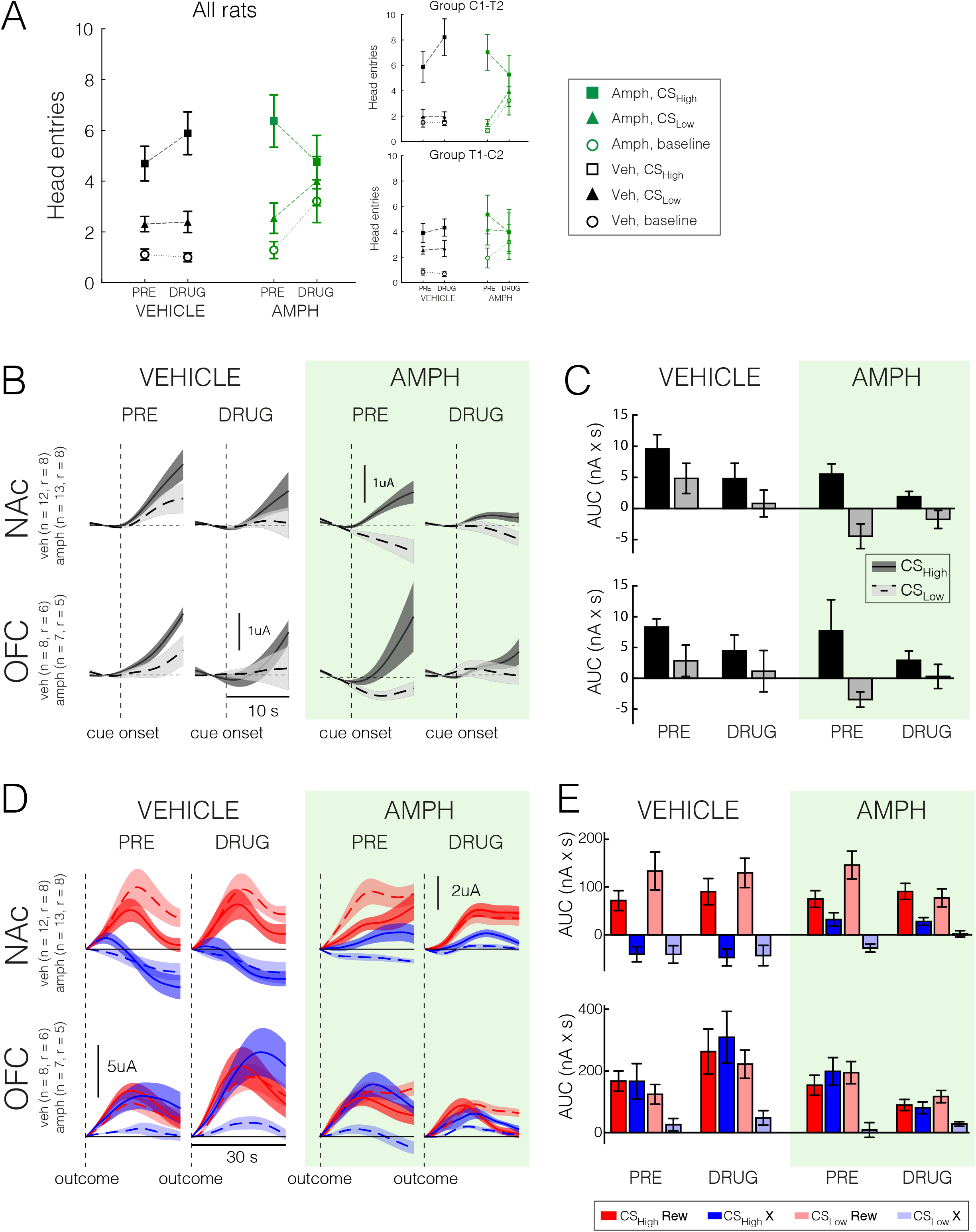
Effect of acute amphetamine administration on cue-elicited behaviour and haemodynamic signals. **A**: Average head entries (mean ± SEM) to the magazine during presentation of each cue or during the pre-Cue baseline period in the session before (“Pre”) and just after drug administration (“Drug”) in the group receiving vehicle or amphetamine (1 mg/kg). **B**. T_O2_ responses time-locked to cue presentation recorded from either NAc (upper panels) or OFC (lower panels) in the pre-drug or drug administration sessions. Note that differences in the pre-drug patterns of signals in the Vehicle and Amphetamine group mainly reflect the unbalanced assignment of included animals from the two cue identity groups. **C**. Average area-under-the-curve responses (mean ± SEM) extracted from 5-10 sec after cue onset for each cue in the two sessions. **D**. T_O2_ responses time-locked to outcome presentation (reward or no reward) after each cue in the two cue identity groups recorded from either NAc (upper panels) or OFC (lower panels) in the pre-drug or drug administration sessions. **E**. Average area-under-the-curve responses (mean ± SEM) extracted from 30 sec following the outcome after each cue in the two sessions.

Administration of amphetamine also had a pronounced but specific effect on T_O2_ responses. During the cue period, the effect in both NAc and OFC mirrored the effect of the drug on behaviour, with amphetamine abolishing the distinction between the average T_O2_ response elicited by presentation of the CS_High_ or CS_Low_ (cue × session × drug: F_1,32_=6.22, p=0.018; CS_High_ v CS_Low_, amphetamine group drug session, p=0.25; all other p<0.006) (Figure 5B,C, S7A). While there were still notable effects of cue identity on signals, follow up comparisons found that there were no reliable differences in T_O2_ responses between the different cue configurations in either group or session (all p>0.27).

Based on the differences between outcome-elicited signals in NAc and OFC observed during training, we split the outcome-elicited T_O2_ data by region. In the NAc, there was a significant cue × session × outcome × drug interaction (F_1,21_=4.82, p=0.04). We focused follow up analyses on each drug group separately without cue identity as a between-subjects’ factor as the NAc electrode exclusion criteria inadvertently biased the distribution of rats assigned to the drug and vehicle groups as a function of cue identity (χ^2^=9.4, df=3, p=0.024) (see Figure S7 for breakdown by counterbalance group).

While vehicle injections caused no changes in NAc signals (no main effect or interaction with session: all F<1.4, p>0.33), amphetamine had a marked influence on outcome-elicited T_O2_ responses, selectively blunting CS_Low_ outcome responses (cue × session × outcome interaction: F_1,12_=22.07, p=0.001; CS_Low_ reward or no reward: pre-drug v drug session, p<0.005; CS_High_, all p>0.22). This meant that, on amphetamine, there was now no reliable distinction between reward-evoked T_O2_ signals based on the preceding cue (p=0.08; all other CS_High_ v CS_Low_ comparisons, p<0.015) (Figure 5D,E). Consistent with this, we also found a significant reduction in the relationship between T_O2_ responses and positive reward prediction errors selectively after amphetamine (comparison of peak effect size on and off drug: session × drug interaction: F_1,21_=8.02, p=0.01; pre-drug v drug session, amphetamine group: p=0.003; vehicle: p=0.66) (note, we did not analyse the negative RPE as this was already largely absent in the pre-drug session in animals showing strong discrimination between the CS_High_ and CS_Low_).

In OFC, there was also a significant change in T_O2_ responses when comparing outcome-elicited signals on the drug session to the pre-drug day (significant cue × session × drug and cue × session × outcome × drug interactions: both F>5.35, p<0.042). In the control group, vehicle injections caused a general increase in all the OFC T_O2_ responses (main effect of testing session: F_1,6_=7.51, p=0.034). By contrast, in the amphetamine group, there was a striking reduction in OFC T_O2_ signals, particularly elicited by CS_High_ cues (main effect of testing session: F_1,5_=5.12, p=0.073; significant cue × session interaction: F_1,21_=29.527, p=0.003). An analysis of the unsigned prediction error signal also resulted in a session × drug interaction (F_1,11_=6.63, p=0.026), though this was driven both by a numeric decrease in the regression weight in the amphetamine group and an increase in the regression weight in the vehicle group.

Taken together, therefore, amphetamine impaired the discriminative influence of CS_High_ and CS_Low_ cues on behaviour and also corrupted the influence of these cue-based predictions on NAc and OFC T_O2_ responses.

## Discussion

The results presented here show that haemodynamic signals in NAc and OFC dynamically track expectation of reward as rats form associations between cues with high or low probability of reward outcome. Importantly, both regions also displayed distinct forms of haemodynamic prediction error signal. NAc signals were shaped by reward expectation and the specific valence of the reward outcome, while in contrast OFC signals did not discriminate the valence of reward outcome, but rather reflected how surprising either reward outcome was. A single dose of amphetamine, sufficient to modulate dopamine activity, caused a loss of discrimination between cues that was evident both behaviourally and in the haemodynamic signatures of reward expectation and prediction error in both regions.

These results extend a previous study of instrumental learning where increases in NAc T_O2_ were observed as rats learned to associate a deterministic cue with receipt of reward upon pressing a lever [35]. The present probabilistic learning study allowed a formal assessment of whether the measured T_O2_ signals displayed features that would categorise them as encoding reward prediction errors (RPEs). To be considered an RPE signal, three cardinal features should be measurable: (i) a *positive* influence of expected reward value on cue-elicited signals (i.e., a greater response to a cue that is thought to predict a higher reward), (ii) a *positive* influence of actual reward delivered (i.e., a greater response when a high value reward is actually delivered compared to when it is omitted) and (iii) a *negative* influence of expected reward value on outcome-elicited signals (i.e., a larger response to reward delivery the less that reward is expected and/or a smaller response to reward omission the more that reward is expected) [43,44]. Using behavioural modelling, all three of those features were evident in NAc T_O2_ signal, consistent with a number of human fMRI studies of reward-guided learning in healthy subjects [43–46]. Although human fMRI studies predominantly use secondary reinforcers such as money to incentivise performance, similar RPE-like activations in NAc are also observed in studies using primary fluid reinforcers in lightly food/water restricted participants, which more closely mimic the means by which rats are motivated to perform the present task [see 47].

While both positive and negative RPE-like T_O2_ signals were evident across the whole learning period, it was clear the influence of each signal changed over time. Both positive and negative RPEs were evident early in learning. However, as discrimination between high and low reward probability cues was learned, negative RPEs had an increasingly negligible influence on NAc haemodynamic signals. Such adaptation has resonance with a previous finding in humans that NAc BOLD signals are not observed for every RPE event, but only those currently relevant to guide future behaviour [46]. The selective involvement of NAc in signalling whether an event is better or worse than expected fits well with the hypothesised roles of the extensive dopaminergic projections to this region, and suggests a fundamental role for NAc in sustaining approach responses to reward-associated cues [48,49].

It has been demonstrated that dopamine release in the core region of the NAc correlates with a RPE and, similar to observed here, dynamically changes over the course of learning [9,10,14,50,51] (though see [52] for a different interpretation). The amperometry electrodes in the current study were largely in caudal parts of ventral NAc, spanning the core and ventral shell regions. Given that the electrodes are estimated to be sensitive to changes in signal over approximately a 400μm sphere around the electrode [53,54], it is plausible that the signals we recorded here could have been influenced by RPE-like patterns of dopamine release [55]. Several recent papers have shown that optogenetic stimulation of dopamine neurons can have widespread influence on forebrain BOLD signals [56–58]. However, direct evidence for this link is currently lacking, and a recent study comparing patterns of BOLD signals with dopamine release in humans found indications of uncoupling between the measures [59]. Therefore, it is also conceivable that the NAc haemodynamic signals we observed here instead reflect afferent input from regions such as medial frontal cortex, where RPE-like signals have also been recorded [44,60,61].

By contrast, OFC T_O2_ signals did not respond to prediction error events in quite the same way as the NAc did and did not meet all three formal criteria to be considered formal correlates of an RPE. While some studies have found RPE-like outcome signals in OFC [62,63], several – including those fMRI studies that have adopted the stringent criteria applied here – have not [e.g., 44,64,65]. Like NAc, OFC T_O2_ signals signalled reward expectations when the cues were presented. This is consistent with previous fMRI and electrophysiological studies suggesting that central or lateral OFC may represent stimulus – reward mappings during cue presentation [e.g., 66,67,68]. However, unlike NAc, the OFC signals measured in the present study tended to *increase* following reward omission as well as after reward delivery and this increase scaled with how surprising the reward omission was. Electrophysiological studies suggest that similar proportions of OFC cells exhibit either positive or negative relationships with value and individual neurons can encode both positive and negative valenced information at outcome [69,70].

These outcome-driven signals did however correlate with an *unsigned* prediction error: how surprising or salient any outcome is based on current expectations. There are an increasing number of studies implicating OFC in modulating salience for the purposes of learning [71–74]. However, given the precise pattern of OFC signals observed in the current study, the OFC T_O2_ responses might reflect the acquired salience of an outcome [e.g., 71,75], which represents both how surprising *and* how rewarding it is. Even though such responses do not signal whether an outcome is better or worse than expected, they are still important to guide the rate of learning or reallocate attention. Single site lesion studies demonstrate a role for OFC, as well as for NAc, in aspects of stimulus-reward learning [76,77]. However, a specific interaction between these regions to support these behaviours must depend upon another mediating region, as there is no direct projection between the two [78].

One unexpected but fortuitous finding was that T_O2_ signals related to magazine responding were strongly influenced by cue identity. By comparing different behavioural models, this effect was best explained by including two additional parameters to a simple reinforcement learning model: (i) a cue salience parameter, which scaled the influence of the RPE on future value estimates as a function of which auditory cue had been presented, and (ii) a cue “arousal” parameter, which was a constant term applied from the start of training. It has long been established that cue salience can be an important determinant of learning rates [e.g., 79,80,81] and this is a standard term in the Rescorla-Wagner and other influential association learning models, capturing the effect that more salient or intense cues are learned about faster and are more readily discriminated than weaker or less salient cues. However, an extra parameter was also needed to account for the fact that in the majority of animals responding to the clicker was greater than the tone at the start of testing irrespective of whether it was assigned to be CS_High_ or CS_Low_. While we do not know the precise reason for this effect and are not aware of this being reported in previous studies, one speculation is that the rats partially generalised the clicker cue to the sound of the pellet dispenser. Regardless of its precise cause, importantly, using model-derived estimates of the value, we were nonetheless able to observe identical neural correlates of reward prediction and prediction error in both cue counterbalanced groups.

Moreover, across both groups, a single “moderate/high” dose of amphetamine (1 mg/kg) [82] at the end of learning, mimicking a dopamine hyperactivity state, was sufficient to impair discriminative behavioural responses to the high and low probability cues and also to disrupt haemodynamic signals in both NAc and OFC. This was not simply a blunt pharmacological influence on neurovascular coupling as the signal change was specific to certain conditions, for instance selectively reducing the NAc reward signals on CS_Low_ trials but leaving the reward and omission responses on CS_High_ trials unaltered.

This may at first appear at odds with previous studies that have shown that this dose of amphetamine can selectively augment responding to reward-predictive cues over neutral cues and enhance phasic dopamine release and neuronal activity in the NAc [36,39] and neuronal activity in OFC [83]. However, there are a number of important differences between these previous studies and the one reported here.

First, in our paradigm, the cues were probabilistically rewarded meaning that both were associated with a certain level of reward expectation and elicit conditioned approach. A previous study has shown that the same dose of amphetamine as used in the current study can disrupt conditional discrimination performance [40]. Second, as discussed above, increased dopamine release may not necessarily map directly onto comparable haemodynamic changes. Indeed, although it would be expected that the dose of amphetamine used in the current experiment would boost phasic dopamine release to reward-predicting cues [36], fMRI studies have tended to observe a blunting of haemodynamic responses to such cues in NAc and frontal cortex following administration of a single dose of amphetamine or methamphetamine, comparable to what we observed in both NAc and OFC [37,84]. In addition, one of these studies [37] reported the loss of RPE encoding in NAc, an effect that was also evident in the current study. Amphetamine is known to increase levels of dopamine – and other monoamines – in a stimulus-independent as well as a stimulus-driven manner. Therefore, the critical factor for appropriate responding is likely to rest on the balance of these two elements in frontal-striatal-limbic circuits, and the disruption of haemodynamic signalling of incentive predictions and prediction errors we recorded may reflect this. From the present data, it is unclear whether amphetamine is directly corrupting calculations of RPEs or is instead primarily disrupting the inputs then used to calculate the RPE such as cue-elicited reward expectation.

The changes in haemodynamic responses observed after amphetamine administration here are also consistent with an increasingly large body of fMRI studies of reward-guided learning in patients displaying symptoms that are believed to arise in part from dysregulated dopamine, such as psychosis and anhedonia (see [16,19,25,85] for reviews). This has raised the possibility that changes in behaviour and brain responses during reward anticipation and reinforcement learning might act as a cross-diagnostic pre-clinical translational biomarker [16,86]. In parallel, there has been increased interest in using these types of finding as foundations for theoretical approaches to link underlying biological dysfunctions to observed symptoms in patients (e.g., [21,87–89]).

However, while there is general consensus about the promise of such approaches, the literature is complicated by the diversity of the disorders and the drug regimens that patients have taken, which makes testing specific causal hypotheses about the relationship between altered brain function and psychiatric symptoms difficult. By contrast, in an animal model, it is possible to have precise control over and measurement of induced changes in neurobiology. Therefore, establishing the feasibility of observing these signatures in a freely behaving rodent, measured using a valid proxy for fMRI, and demonstrating how clinically relevant pharmacological perturbations affect these responses, may be an important step to bridge the gaps in our understanding. This foundation, if combined with other causal manipulations (such as pharmacological or genetic animal models relevant to psychiatric disorders) and more sophisticated behavioural tasks that allow us to take into account different learning strategies [e.g., 90,91] and value parameters [7,92], could therefore provide new opportunities for understanding how dysfunctional neurotransmission are reflected as changes in haemodynamic signatures and how both relate to behavioural performance.

## Acknowledgements

MEW, MT, HM and GG conceived the project, MEW, GG, FR, and JF designed the experiment, JF and FG collected the data, JF performed the surgeries, EW, SS, JH, and MEW analysed the data, MEW prepared the manuscript with input from the other authors. We would like to thank David Bannerman for valuable advice throughout the project, Mike Conway for assistance with surgeries, and Thomas Akam, Miriam Klein-Flugge, Stephen McHugh and Marios Panayi for discussions about the modelling, interpretation and analyses. The study is dedicated to the memory of one of the authors, Soon-Lim Shin, a key contributor to the project who sadly passed away to cancer in 2017.

