## Supplementary Material for "Amphetamine disrupts haemodynamic correlates of prediction errors in nucleus accumbens and orbitofrontal cortex"

### Supporting Information

#### Methods

##### *Animals*

Adult male Sprague Dawley rats (Charles River, UK) were used in the present studies (*n*=36 at start). The sample size was chosen based on previously reported robust $O_2$ signal differences related to cue presentation in reward-guided learning tasks [1,2]. 4 animals did not contribute to the behavioural dataset owing to poor $O_2$ calibration responses and the data from an additional 2 animals could not be included owing to a computer error, resulting in a total of 30 rats providing behavioural data. Before surgery, animals (400-450g) were housed in standard housing conditions (four per cage, 7:00 A.M. to 7:00 P.M. light phase, controlled temperature and humidity, *ad libitum* water). After surgery, animals were singly housed in the same environment. All animals were kept for a period of 7 d before any behavioural procedure started. During this time, rats were acclimated to the food restriction regime (i.e., maintained at no less than 85% of their free-feeding weight relative to their normal growth curve) and were handled regularly. For environmental enrichment, home cages were supplied with wooden blocks and nesting materials.

##### *Electrode construction*

Carbon paste electrodes (CPEs) were constructed from 8T (200 μm bare diameter, 270 μm coated diameter) Teflon-coated silver wire (Advent Research Materials). The Teflon insulation was slid along the wire to create a ~2 mm deep cavity, which was packed with carbon paste. Carbon paste was prepared by thoroughly mixing 7.1 g of carbon graphite powder and 2.5 ml of silicone oil (both from Sigma-Aldrich, UK). Reference and auxiliary electrodes were also prepared from 8T Teflon-coated silver wire by removing the Teflon from the tip. All electrodes were soldered to gold connectors, which were cemented into six-pin plastic sockets (both from Plastic One, UK) during surgery.

##### *Electrode calibration*

Before implantation, all CPEs were calibrated *in vitro* in a standard three electrode glass electrochemical cell (C3 cell stand, BASi), with an Ag/AgCl reference electrode and a BASi platinum auxiliary electrode. Calibrations were performed in a 15ml phosphate buffer saline solution, pH 7.4, saturated with nitrogen (N2) gas, atmospheric air (from a RENA air pump), or pure O2 at room temperature providing a 3-point calibration of known concentrations of 0 µM (N2 saturated), 240 µM (air saturated), and 1260 µM (O2 saturated) oxygen. CPEs were chosen for implantation if their calibration curves were linear and the measured oxygen values from the saturated solutions were not greatly different from those expected (least square linear regression, $R^2 \geq 0.98$).

*Surgery*

Surgery was performed in experimentally naïve rats, prior to any training or testing. Animals were placed in vaporization chambers and anesthetized with 5% isofluorane (2 L/min $O_2$) and maintained on 1–3% isofluorane (2 L/min $O_2$) for the rest of the procedure. CPEs were implanted in the nucleus accumbens (NAc) [from bregma: anteroposterior (AP), +1.4 mm; mediolateral (ML), ±1.3 mm; dorsoventral (DV), –6.4mm] and the orbitofrontal cortex (OFC) [from bregma: AP,+3.7 mm; ML, ±2.4 mm; DV, –5.2 mm)]. The reference electrode was inserted into the posterior cortex to a depth of 2 mm and secured with cement. The auxiliary electrode was wrapped round a skull screw positioned over posterior cortex. After all electrodes were cemented into place, the gold sockets of the electrodes were inserted into a six-pin plastic socket. All animals were administered Rimadyl (carprofen; 5 mg/kg, s.c.) both presurgically and postsurgically, and were allowed to recover in thermostatically controlled cages.

*In vivo signal validation*

To demonstrate the oxygen sensitivity and reliability of the electrodes *in vivo*, mild hyperoxia and hypoxia were induced by applying gaseous $O_2$ (BOC Medical) or $N_2$ (BOC Gases), respectively, to the snout of the animal before and after each experiment. Polyurethane tubing, connected to the appropriate gas cylinder, was held ~2 cm from the snout and the gas delivered for either 60 s ($O_2$) or 30 s ($N_2$) at a flow rate of 1L/min. Electrodes were considered acceptable for analysis if they did not show any increase in response following two $N_2$ inhalation challenges, and a reproducible increase to two $O_2$ inhalation challenges.

*O2 amperometry data recording*

Rats were connected to a four-channel potentiostat (Biostat, ACM Instruments) through a six-pin socket (Plastics One) via a flexible screened six-core cable (Plastics One). A PowerLab 8/30 Data Acquisition System was used for analogue/digital conversion, and data were collected on a PC running Chart version 5 software (AD Instruments). The O2 signal was recorded at a sample rate of 200 Hz. For all test sessions where an amperometric signal was recorded, animals were tethered and a constant potential (–650 mV) was applied for the duration of the session.

*Behavioural experiments*

*Apparatus.* Operant boxes housed in sound- and light-attenuating chambers were used (Med Associates). Each chamber contained a house light (100 mA; ENV 215M, Med Associates) and two retractable levers. The levers were located on either side of a recessed magazine where food pellets (45 mg; Formula P, Noyes) were delivered from an automatic pellet dispenser. Auditory signals were produced by a tone generator located on the wall opposite to the food magazine. Experimental sessions were controlled and data recorded using in-house programs written with MedPC-IV software (Med Associates), and data were prepared for analysis using an in-house Excel macro designed for each experiment.

*Probabilistic Pavlovian conditioning task*

A schematic of the experimental design is shown in Figure S1. Initially, rats received 1-2 initial habituation sessions, where they learned to nose poke for food reward under a Variable Interval (VI) 15s schedule. They then progressed to perform a probabilistic Pavlovian learning task. Each trial consisted of a 10s presentation of one of two auditory cues (3 kHz pure tone at 77dB or 100 Hz clicker at 76 dB) followed immediately by either delivery or omission of reward (4 x 45 mg sucrose food pellets) according to a probabilistic schedule. One of the auditory cues was assigned as the $CS_{High}$ and was followed by reward delivery on 75% of trials, the other the $CS_{Low}$ rewarded on 25% of trials, counterbalanced across animals. Each session started with the illumination of the house light and consisted of a total of 56 cue presentations, with an average inter-trial interval of 45s (range 30-60s). Standard training took place over 9 sessions and session 10 consisted of the drug challenge.

*Pharmacological manipulations*

D-amphetamine sulphate (Sigma,UK) was dissolved in 5% (w/v) glucose solution, and pH adjusted towards neutral with the dropwise addition of 1M NaOH as necessary. Amphetamine was dosed at 1mg/kg (free weight) via the intraperitoneal route, 30 minutes prior to the session. Half the animals received amphetamine and the other half vehicle. This corresponds well with critical reviews of published amphetamine studies [3]. 1mg/kg represents a moderate dose of amphetamine that is behaviourally active, yet does not cause too much motor stereotypy that could potentially confound the ability to engage in operant behaviours. This dose will also promote the release of dopamine in the nucleus accumbens, among other regions [e.g., 4]. In humans, positive and/or psychotic-like symptoms would become emergent around this dose range [5,6]. Drug treatment was blinded during dosing.

*Behavioural modelling*

Head entries during the 10 seconds cue presentation were modelled using variations of a Rescorla-Wagner model (Rescorla & Wagner 1972). The value attributed to each cue was updated trial-by-trial (t) as such:

$$v_{cue}(t) = v_{cue}(t-1) + \alpha(R - v_{cue}(t-1))$$

, where $v_{cue}$ is the value for a given cue, R is the receipt (=1) or omission (=0) of reward and α (restricted between 0 and 1) is the reward learning rate.

In order to test whether rats were learning faster from rewarded trials than from non-rewarded trials, we tested models using two different learning rates for rewarded and non-rewarded trials.

$$v_{cue}(t_{rew}) = v_{cue}(t_{rew} - 1) + \alpha_{rew}\big(R - v_{cue}(t_{rew} - 1)\big)$$
$$v_{cue}(t_{no\ rew}) = v_{cue}(t_{no\ rew} - 1) + \alpha_{no\ rew}(R - v_{cue}(t_{no\ rew} - 1))$$

In the other models, described as "single learning rate", all trials used the same learning rate α.

In order to account for potential differences in the salience of the cues influencing the speed of learning, we tested models including a cue salience learning rate. In these models, the learning rate was multiplied by a cue salience term β for trials where a given cue was presented so that:

$$v_{tone}(t) = v_{tone}(t-1) + \alpha\beta_{tone}\big(R - v_{tone}(t-1)\big)$$
$$v_{clicker}(t) = v_{clicker}(t-1) + \alpha\beta_{clicker}\big(R - v_{clicker}(t-1)\big)$$

The values for each cues were then scaled up into head entries units using a parameter *k,* fitted individually for each subject.

$$Predicted\ Head\ Entries(t) = k.v_{cue}(t)$$

To account from differences in responding observed immediately at the start of training, we tested models that included a constant, *C*, to account for cue-induced unconditioned magazine responding. This parameter could either account for a general increases in responding upon presentation of either cue ($C_{general}$, same value for both cues) or a specific differential increase to one cue compared to the other ($C_{clicker}$ and $C_{tone}$). As it is independent of learning, it is present from trial 1.

To account for trial-to-trial fluctuations of head entries explained by fluctuations in non-specific factors such as alertness and satiety, we tested some models in which we added for each trial a smoothed average of the head entries during the baseline of the preceding trials. More precisely, it consisted in an exponential moving average of the preceding baselines, weighed by an excitement factor *E* that regulated, for each rat, the contribution of the baseline to predicted head entries.

We then combined the different elements of the model to calculate the predicted head entries as such:

$$Predicted\ Head\ Entries(t) = k.v_{cue}(t)\ +\ C\ + (E.baseline\ head\ entries)$$

We then calculated the likelihood of each observation as the Poisson probability to observe the real data given the predicted data. The log-likelihood of a model was calculated as the sum of log-likelihood of all trials from all rats. We selected the best-fit parameters that maximised the log-likelihood for each rat, within a constraint range (*α, β and E*: 0-1, C: 0-10, k: 0-100). In order to do so, we ran fmincon, a MATLAB function that minimises constrained functions, on the negative log-likelihood, starting with random initial parameters at least 10 times, and until the 4 best results gave a log-likelihood that was less than 0.1% different from the best likelihood, suggesting a global minimum was found.

To compare the models, we used the Bayesian information criterion (BIC), which penalises the likelihood of a model by the number of parameters and the natural logarithm of the number of data points. The model with the lowest BIC score was deemed to give a better fit of the data. Note, however, that the patterns of results remained unchanged if we used any of the 3 models that fitted best for a number of individual rats (Fig. 1d: models a, c, f) or even if we used a standard Rescorla-Wagner-type learning model (Fig 1d: model s).

*Data analysis*

*Behaviour*

We initially analysed the behaviour in all 30 animals that had performed the task, irrespective of whether their $O_2$ data met our inclusion criteria. To determine whether the animals were learning cue-reward associations, we submitted the average number of head entries into the food magazine during presentation during either the $CS_{High}$ or $CS_{Low}$ cues to a repeated measures ANOVA, with cue and session as within-subjects factors and cue identity (whether the clicker or tone was assigned as the $CS_{High}$ cue) as a between-subjects factor. Owing to a computer error, the data from session 7 in 14/30 rats and also from session 4 in 1 rat could not be included. Therefore, this analysis included 29 rats and the factor of day had 8 levels (sessions 1-6, 8, 9). We also repeated this analysis (i) by using only the subset of 20 rats that had also provided useable $O_2$ signals and (ii) by averaging the sessions into 3 blocks of 3 sessions (thereby including all 30 rats and incorporating the data from session 7 in those animals that provided useable data) and replicated all findings. In cases where the assumption of sphericity was not met, we applied a Huynh-Feld correction and degrees of freedom were updated accordingly. For the amphetamine challenge, we compared the pre-drug day (day 9 of the probabilistic learning) with the drug day using a similar approach, including within-subjects factors for cue and session and cue identity and drug group (amphetamine or saline) as between-subjects factors. Here, we separately analysed baseline magazine responding during the 10s period just before cue presentation as well as cue-elicited responses.

*Amperometry:*

The $T_{O2}$ signals were first low-pass filtered below 0.1 Hz using a biquad Butterworth filter. All filtering was performed in both directions to eliminate phase-shifting of the data. The signals were then down-sampled to 2 Hz, by averaging 0.5 s of data around every 0.5 s time stamp.

As a decrease in the recorded current reflects an increase in oxygen concentration, the data were inverted for presentation.

We first checked for channels that either had no meaningful signal or had excessive noise. To detect flat channels, we measured the root-mean-square (RMS) of the raw data 25 minutes after of the start of recording (to allow the signal to stabilise) and averaged the data over 9 days. Channels with an average RMS value lower than $5x10^{-4}$ were excluded. To detect noisy channels, we measured both the maximum range of the processed data for each trial (i.e., the difference between the maximum and minimum value) and the ratio between the remaining power between 0.1 and 1 Hz after filtering the power between 0 and 0.1 Hz. Again, the data over 9 days were averaged together. Channels were excluded if they had an average maximum range over 35 and/or a power ratio over 0.0065 (Figure S1).

To understand the relationship between the $T_{O2}$ signals and behaviour, we performed two sets of complementary analyses: (i) model free analyses, where we investigated the average signals in NAc and OFC over the course of learning and after amphetamine administration, and (ii) model based analyses where we regressed the took the same signals against estimates from our computational model.

For the model free analyses, we compared the signal amplitude in response to specific behavioural time points by computing the area under the processed oxygen signal. For responses to cue presentation, the signal was zeroed at cue onset, and the area under the curve was calculated over 10 seconds from cue onset. For responses to outcome, the signal was zeroed at outcome and the area under the curve was calculated over 30 seconds from outcome. This allowed us to examine the relative change in $T_{O2}$ responses to significant events in the task. We subjected the average aligned $T_{O2}$ responses to a repeated measures ANOVA, with within subjects factors of cue ($CS_{High}$, $CS_{Low}$) and training stage (averaged over 3-days into 'early', 'mid' and 'late' stages) and between subjects factors of cue identity (CL1-T2, T1-CL2) and brain region (NAc, OFC).

For the model based analyses, regression coefficients were estimated for each animal at each time point (every 0.5 s) in a 20s window spanning the cue period (10s before cue onset and the 10s of cue presentation, $T_{O2}$ responses zeroed at cue onset) or a 30s window after reward delivery or omission ($T_{O2}$ responses zeroed at outcome). We fitted a generalised linear model (GLM) with a constant term and a set of regressors. These could include: **expected value** (a combination of learned cue value and unconditioned cue-elicited responding: $EV_{cue} =$

$v_{cue} + \frac{C\,cue}{k}$); **outcome** (reward 0.5, no reward -0.5); **RPE** ($R - EV_{cue}$), with the positive RPE regressor built as from positive values of the RPE and set to 0 for the negative RPE and likewise for the negative RPE regressor; **unsigned PE**, which was the absolute value of the RPE. We also included regressors of no interest to account for fluctuations in $T_{O2}$ responses related to satiety (cumulative rewards received per session); outcome on the previous trial; and average recent baseline head entries. All regressors, apart from positive / negative RPEs, were mean-centred. Regression coefficients in each animal were averaged and significance was assessed by comparing these values against a population of 1000 coefficients obtained by randomly permuting the values of the regressors. Significance was determined if the real data exceeding the maximum or minimum of the permuted population of coefficients (i.e., $p < 0.001$ uncorrected). To compare regression coefficients with behaviour or examine changes in regression coefficients before and after drug administration, the peak coefficient was extracted and analysed. To examine the effect of amphetamine on prediction error signals in these sessions, we used fixed idealised values rather than subject-specific estimates (0.5 and 1 for the positive RPE, and –1 and –0.5 for the negative RPE, of CS1 and CS2 respectively).

*Histology*

Animals were deeply anaesthetized with pentobarbital and perfused transcardially with 0.9% (w/v) saline followed by 10% (w/v) buffered paraformaldehyde solution. Brains were removed, placed in formalin, and shipped for histological processing (Covance, Greenfield, IN). The brains were trimmed to include areas of interest, embedded in paraffin, and step-sectioned at 200 µm (4µm per section). Hematoxylin and eosin-staining was used to evaluate electrode placement with reference to a standard rat brain atlas (Paxinos and Watson, 2009). All inaccurate placements were excluded from analyses of the amperometric data.

#### Supplementary Figure Legends

##### Figure S1. Schematic of the surgery, training and testing schedule

**Figure S2. Behavioural performance of the subset of rats (n=20) from which at least 1 channel of oxygen data was analysed.** Upper panel, all rats contributing amperometric data; lower panels, same data split into the two counterbalance groups.

**Figure S3. Electrode inclusion criteria. A.** Histogram of the average root mean square (RMS) of the raw data over 9 days in each of the recorded channels. Channels with classified as "flat" and excluded if they had an average RMS value lower than $5x10^{-4}$. **B.** Scatterplot of the maximum range of the processed data for each trial plotted against the ratio between the remaining power between 0.1 and 1 Hz after filtering the power between 0 and 0.1 Hz for each channel, averaged together over the 9 days. Channels were excluded if they had an average maximum range over 35 (blue vertical line) and/or a power ratio over 0.0065 (red horizontal line).

**Figure S4. Comparing the influence of model-derived value signals and magazine responding on haemodynamic signals. A.** Average effect sizes in NAc (left panel) and OFC (right panel) from separate general linear models relating $T_{O2}$ responses in the 10s before and during CS presentation either to (a) trial-by-trial estimates of the expected value associated with each cue or (b) trial-by-trial magazine head entries during CS presentation. The effect size for expected value was significantly greater than for magazine head entries in both regions (p = 0.024 and p = 0.041 for NAc and OFC respectively). **B.** Same as A, except comparing $T_{O2}$ responses in the 10s before and during CS presentation to the number of head entries in the 10s pre-cue baseline period.

**Figure S5. Influence of negative RPEs on NAc responses change over training. A.** Average effect sizes in NAc from a general linear model relating $T_{O2}$ responses to trial-by-trial estimates of positive or negative reward prediction errors reward or no reward events respectively in the two groups over training. **B.** Relationship between peak effect size for a positive or negative RPE regressor recorded early (sessions 1-3), mid (sessions 4-6) or late (sessions 7-9) in training plotted against a CS discrimination index (the difference in cue-elicited responding to the $CS_{High}$ or $CS_{Low}$).

**Figure S6. Influence of unsigned PEs on OFC responses.** Average effect sizes in OFC from a general linear model relating $T_{O2}$ responses to trial-by-trial estimates of unsigned prediction errors. Main plots include all animals, insets show the analyses divided up into the two cue identity groups.

**Figure S7. Effect of acute amphetamine administration on haemodynamic signals divided by counterbalance group.** **A**. $T_{O2}$ responses time-locked to cue presentation recorded from either NAc (upper panels) or OFC (lower panels) in the pre-drug or drug administration sessions. **B**. $T_{O2}$ responses time-locked to outcome presentation (reward or no reward) after each cue in the two cue identity groups recorded from either NAc (upper panels) or OFC (lower panels) in the pre-drug or drug administration sessions.

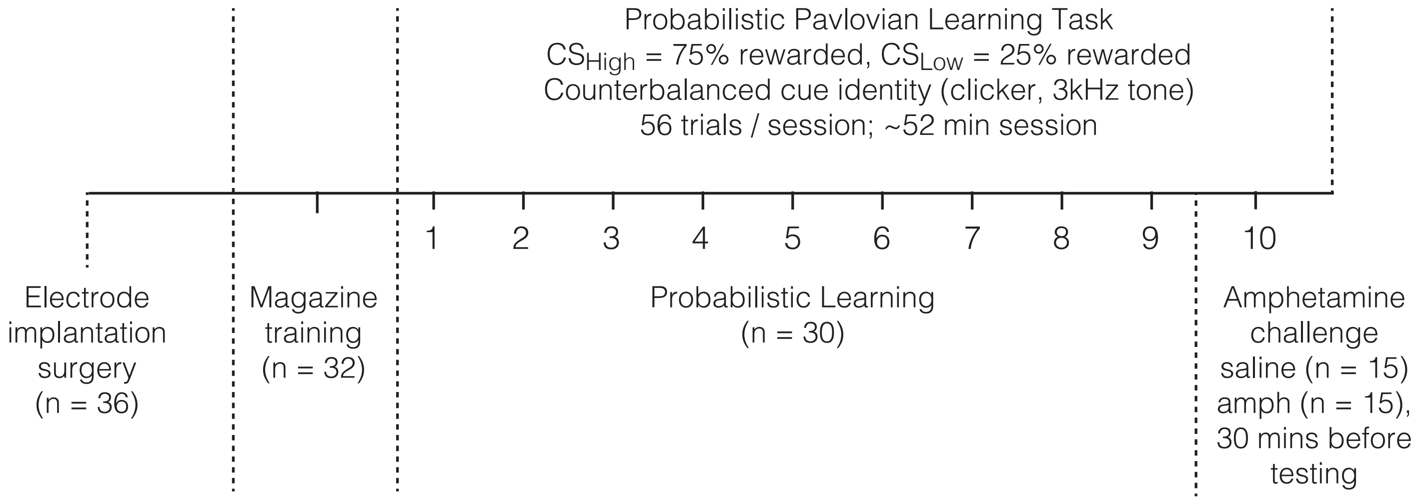
Probabilistic Pavlovian Learning Task
$CS_{High}$ = 75% rewarded, $CS_{Low}$ = 25% rewarded
Counterbalanced cue identity (clicker, 3kHz tone)
56 trials / session; ~52 min session
1
2
3
4
5
6
7
8
9
10
Electrode implantation surgery (n = 36)
Magazine training (n = 32)
Probabilistic Learning (n = 30)
Amphetamine challenge saline (n = 15) amph (n = 15), 30 mins before testing

Figure S1

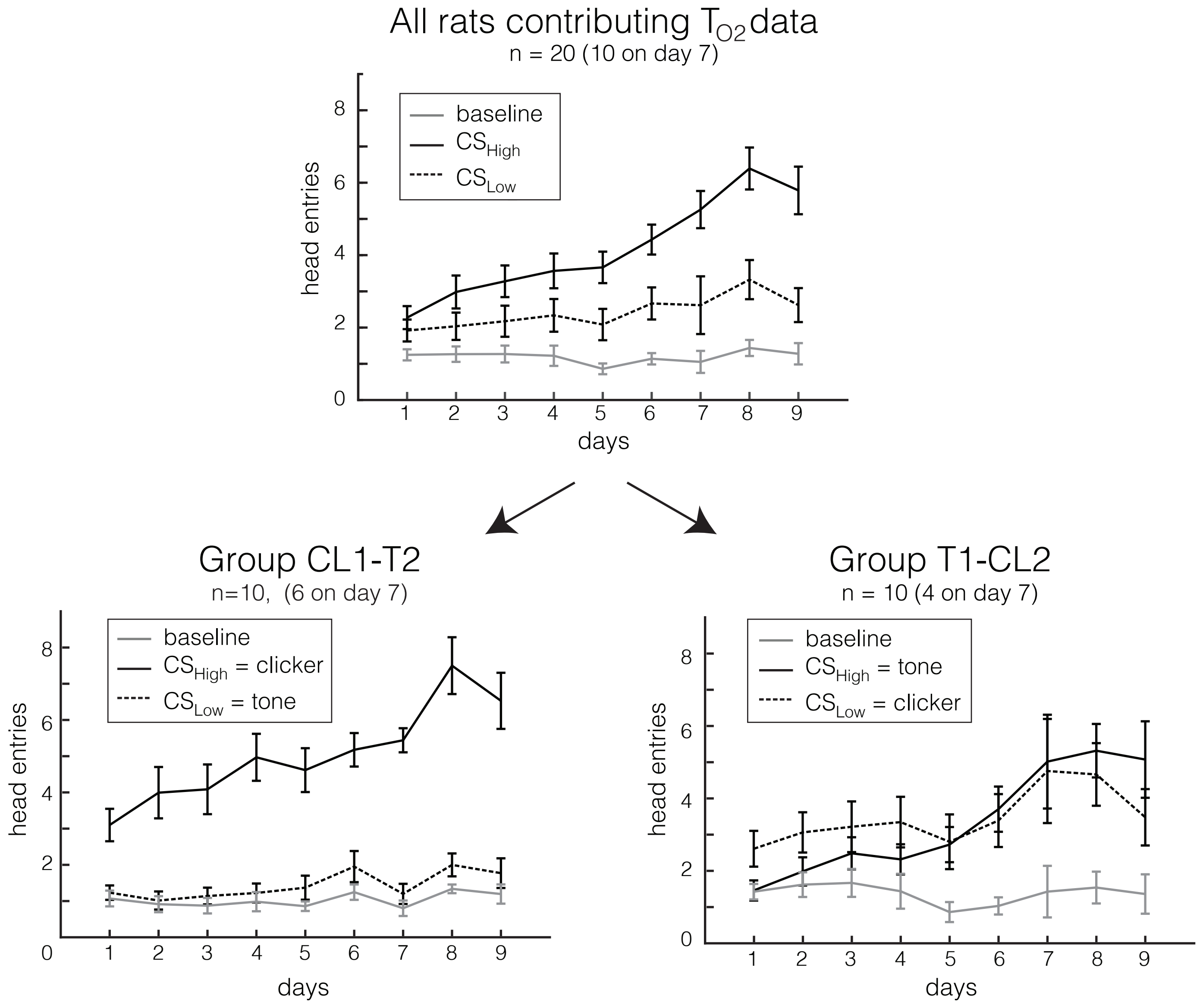

All rats contributing $T_{O2}$ data
n = 20 (10 on day 7)
baseline
$CS_{High}$
$CS_{Low}$
head entries
days
0
2
4
6
8
1
2
3
4
5
6
7
8
9
Group CL1-T2
n=10, (6 on day 7)
baseline
$CS_{High}$ = clicker
$CS_{Low}$ = tone
head entries
days
Group T1-CL2
n = 10 (4 on day 7)
baseline
$CS_{High}$ = tone
$CS_{Low}$ = clicker
head entries
days


Figure S2

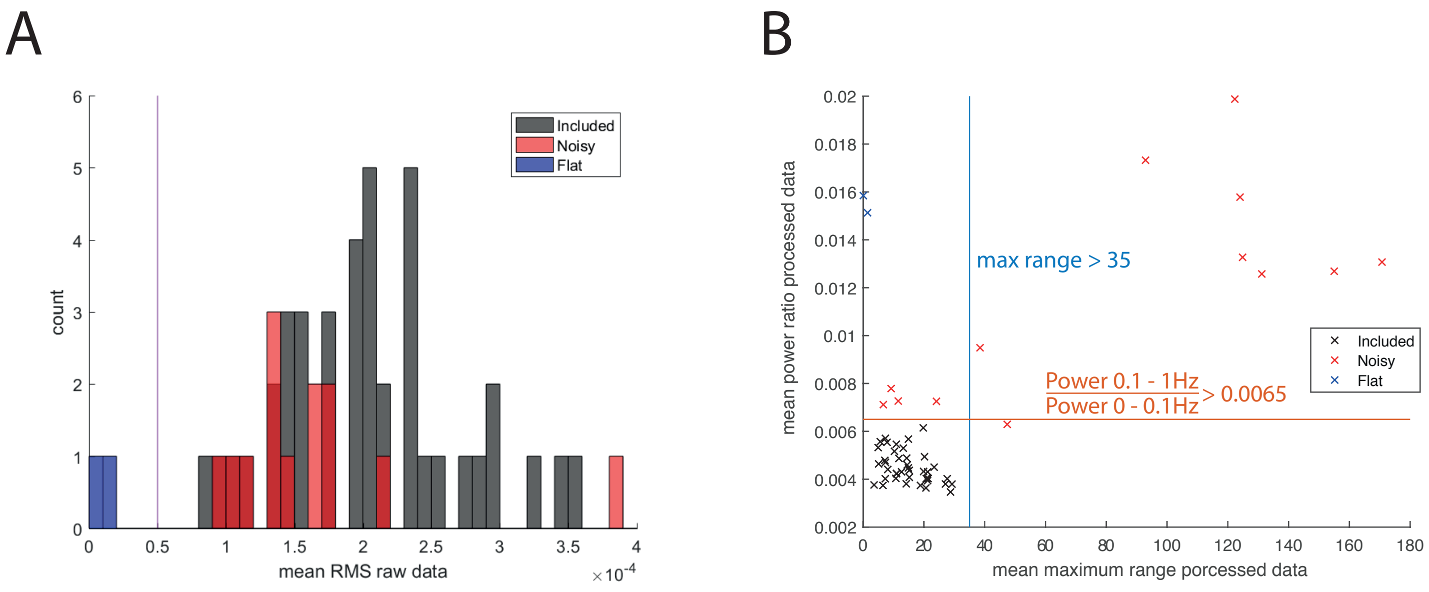
A
B
count
Included
Noisy
Flat
mean RMS raw data
×10^-4
mean power ratio processed data
max range > 35
Power 0.1 - 1Hz
Power 0 - 0.1Hz
> 0.0065
mean maximum range porcessed data

Figure S3

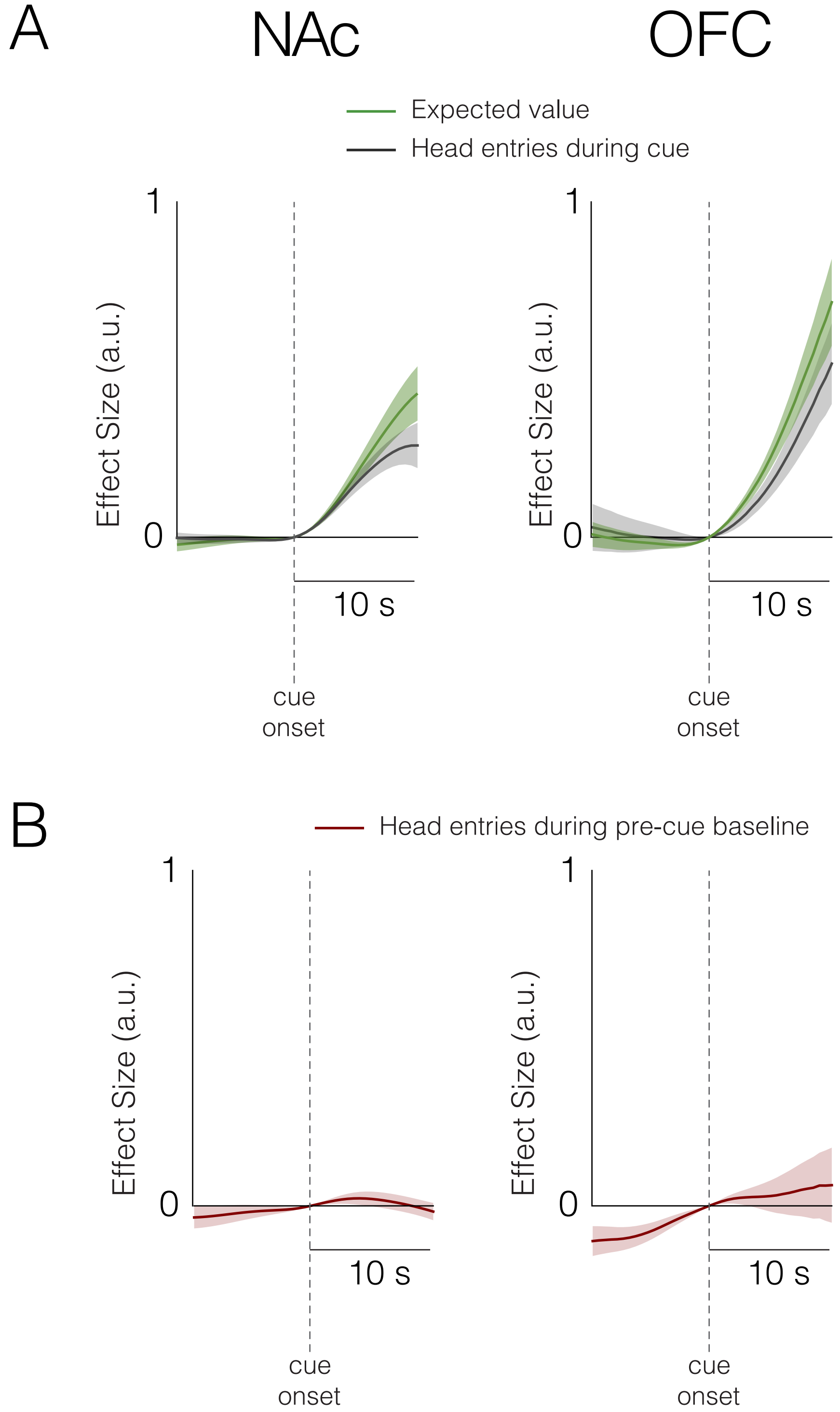
A
NAc
OFC
Expected value
Head entries during cue
1
0
Effect Size (a.u.)
10 s
cue
onset
B
Head entries during pre-cue baseline
1
0
Effect Size (a.u.)
10 s
cue
onset

Figure S4

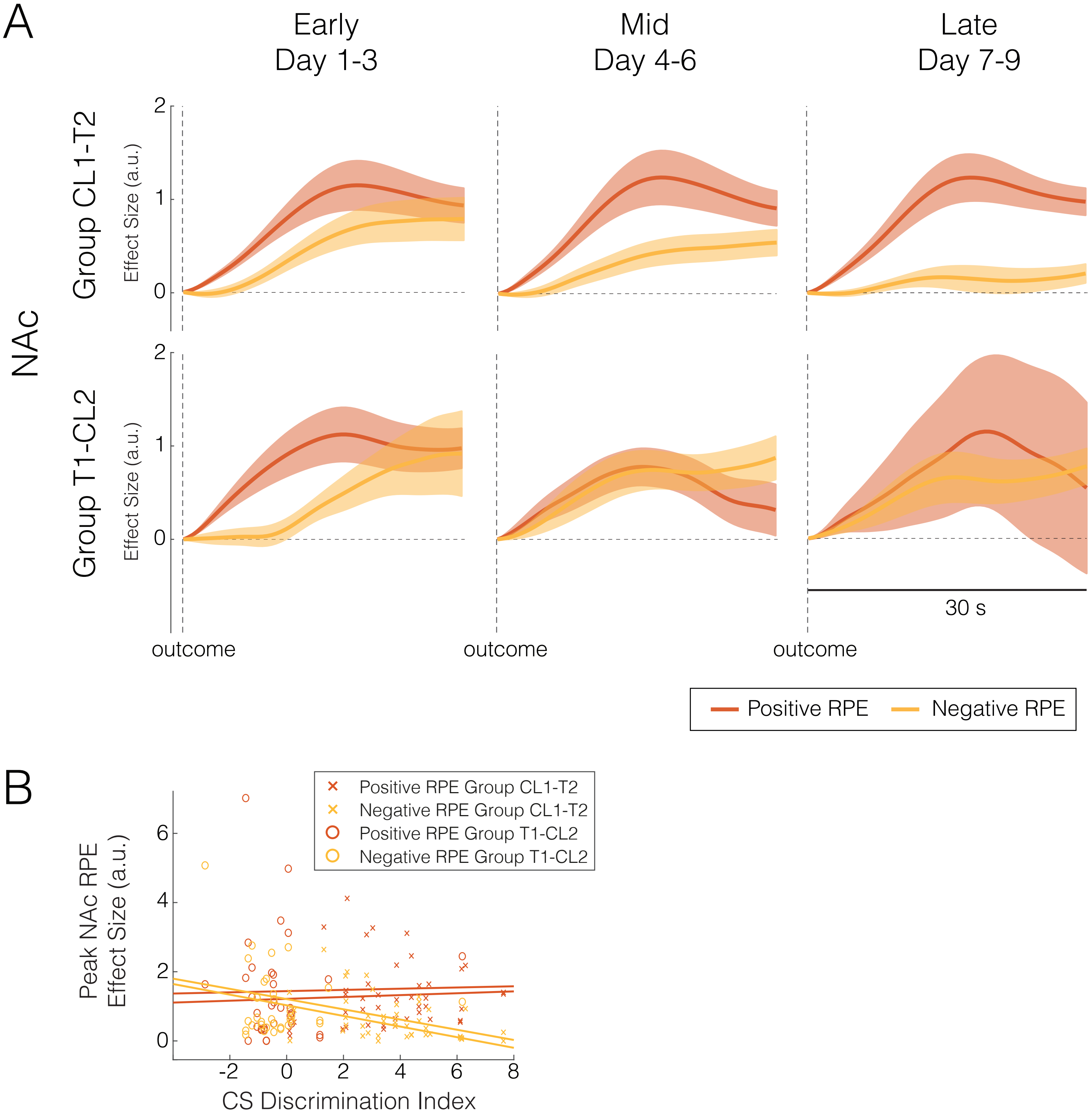
A
Early
Day 1-3
Mid
Day 4-6
Late
Day 7-9
NAc
Group CL1-T2
Group T1-CL2
Effect Size (a.u.)
2
1
0
outcome
30 s
Positive RPE
Negative RPE
B
Positive RPE Group CL1-T2
Negative RPE Group CL1-T2
Positive RPE Group T1-CL2
Negative RPE Group T1-CL2
Peak NAc RPE
Effect Size (a.u.)
6
4
2
0
-2
8
CS Discrimination Index

Figure S5

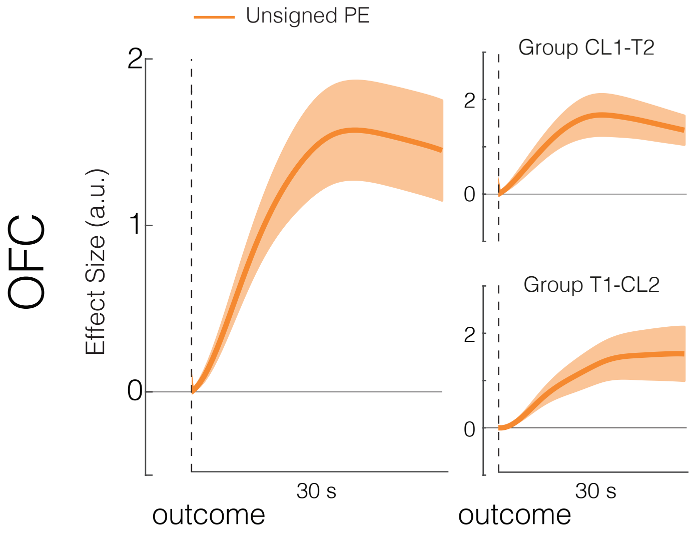

Unsigned PE
OFC
Effect Size (a.u.)
2
1
0
30 s
outcome
Group CL1-T2
2
0
Group T1-CL2
2
0
30 s
outcome


Figure S6

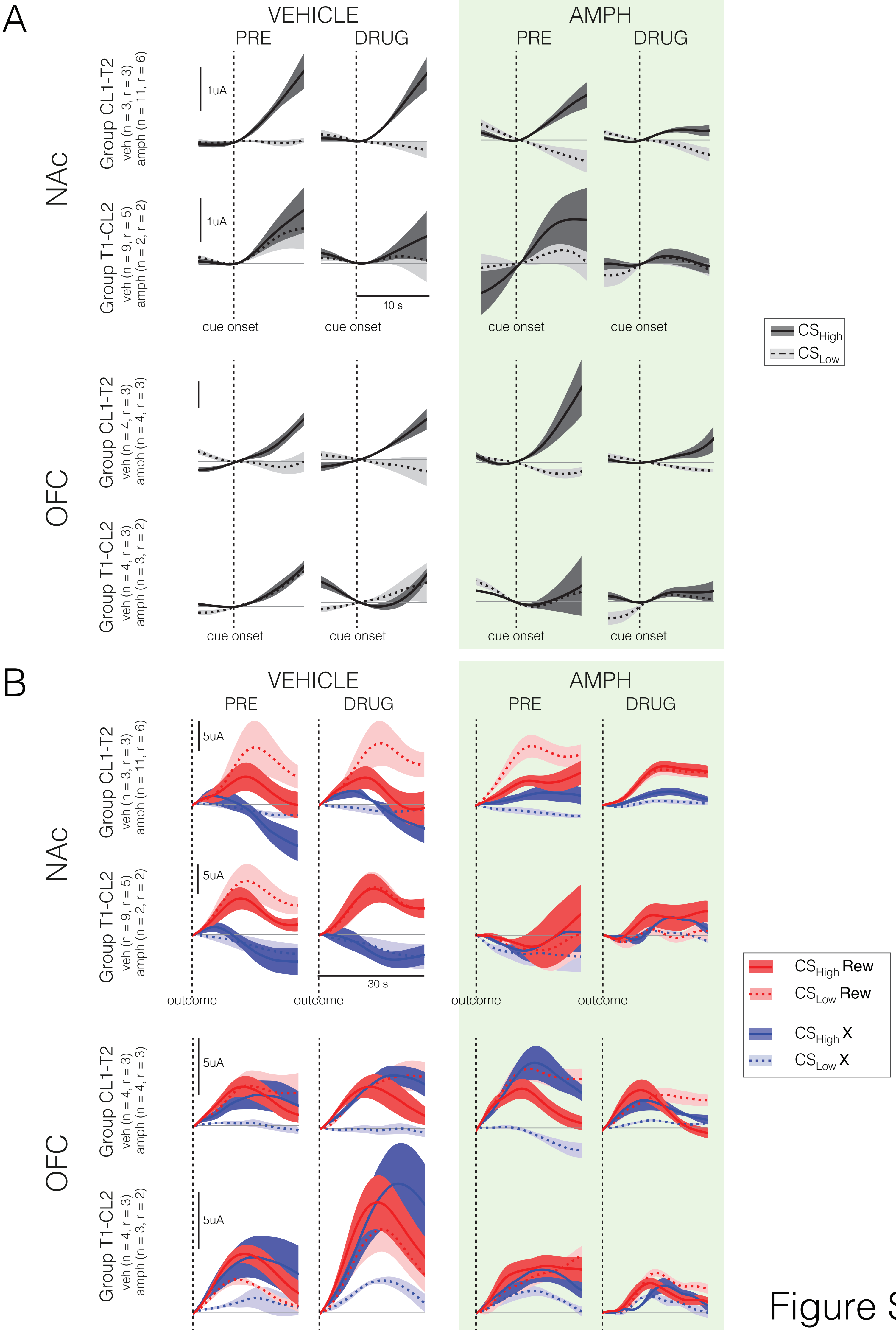
A
VEHICLE
PRE
DRUG
AMPH
PRE
DRUG
NAc
Group CL1-T2
veh (n = 3, r = 3)
amph (n = 11, r = 6)
1uA
Group T1-CL2
veh (n = 9, r = 5)
amph (n = 2, r = 2)
1uA
10 s
cue onset
cue onset
cue onset
cue onset
CSHigh
CSLow
OFC
Group CL1-T2
veh (n = 4, r = 3)
amph (n = 4, r = 3)
Group T1-CL2
veh (n = 4, r = 3)
amph (n = 3, r = 2)
cue onset
cue onset
cue onset
cue onset
B
VEHICLE
PRE
DRUG
AMPH
PRE
DRUG
NAc
Group CL1-T2
veh (n = 3, r = 3)
amph (n = 11, r = 6)
5uA
Group T1-CL2
veh (n = 9, r = 5)
amph (n = 2, r = 2)
5uA
30 s
outcome
outcome
outcome
outcome
CSHigh Rew
CSLow Rew
CSHigh X
CSLow X
OFC
Group CL1-T2
veh (n = 4, r = 3)
amph (n = 4, r = 3)
5uA
Group T1-CL2
veh (n = 4, r = 3)
amph (n = 3, r = 2)
5uA

Figure S7
